# Variable Mutations at the p53-R273 Oncogenic Hotspot Position Leads to Altered Properties

**DOI:** 10.1101/684407

**Authors:** A Garg, J P Hazra, M K Sannigrahi, S Rakshit, S Sinha

## Abstract

Mutations in p53 protein, especially in the DNA binding domain is one of the major hallmarks of cancer. The R273 position is a DNA contact position and has several oncogenic variants. Surprisingly, cancer patients carrying different mutant-variants of R273 in p53 have different survival rates indicating that the DNA contact inhibition may not be the sole reason for reduced survival with R273 variants. Here, we probed the structural properties of three major oncogenic variants of the R273: ([R273L], [R273H], and [R273C])p53. Using a series of biophysical, biochemical and theoretical simulation studies, we observe that these oncogenic variants of the p53 not only suffer a loss in DNA binding, but also show distinct structural stabilty, aggregation and toxicity profiles. [R273C]p53 shows maximum amyloidogenicity while [R273L]p53 shows maximum aggregation. Further probe in the aggregation mechanism show that [R273C]p53 aggregation is disulphide mediated whereas hydrophobic interactions dominate self-assembly in [R273L]p53. MD simulation studies clearly show that α-helical intermediates are observed in [R273C]p53 whereas β-sheets are observed for [R273L]p53. Our study indicates that each of the R273 variant has its own distinct property of stability and self-assembly, the molecular basis of which, may lead to different types of cancer pathogenesis *in vivo*. These studies will aid the design of therapeutic strategies for cancer using residue specific or process specific protein aggregation as target.

**Statement of significance:** The present work stems from an interesting observation that genetic mutations that results in switching of one amino acid to different variants at the same codon show different cancer cell progression. We are trying to understand the molecular reason behind the different gain-of-function opted by these variants. With the help of biophysical and biochemical experiments, and computational studies we have observed that the different thermal stability, unique mechanism of unfolding and self-assembly might be one of the crucial parameters for their different oncogenic effect. These studies thus call for the need of developing therapeutic strategies that consider the resultant mutant-variant as a target rather than mutation position. This is an important lead towards the understanding of cancer.

## Introduction

Abnormal protein behavior is the key cause to several clinical conditions. The anomaly in the protein behavior can be due to mutations, misfolding of the protein, or non-clearance of the protein by degradation pathways leading to accumulation and aggregation. Protein aggregation is linked to several neurodegenerative diseases like amyloid beta aggregation in Alzheimer’s disease, aggregates of α-synuclein in Parkinson’s disease(1-6). Over the last few decades, a good amount of work suggests that the misfolding and aggregation of the p53 protein is a pathological condition in cancer (7-9). The p53 protein, central to the human physiology, involved in maintaining genomic stability by inducing cell-cycle arrest, senescence and apoptosis(10-13), is found to be frequently mutated in cancer and majority of the mutations are present in the DNA binding domain of p53(14-16). Selective mutations in this central DNA binding domain are more frequent in tumor and are thought to be an important factor for their uncontrolled growth(17, 18). It is also reported that a few of these mutations alter the structure of the protein that lead to its misfolding and aggregation (19-21). R273 and R248 are two such hotspot DNA contact mutations which show maximum frequency of alteration in cancer(22, 23). R273 is one of previously defined DNA-contact position, the mutation of which is shown to have the most severe impact on cancer patients. The R273 position has the highest frequency of mutation in the cancers of the brain, ovary, pancreas, and liver (24-26) (Figure 1, S1a, Table S1). The R273 residue is mutated to different amino acids in different types of cancer. The three most common variants of R273 mutations observed in tumors are [R273H]p53, [R273L]p53 and [R273C]p53. Previous report shows that [R273H] and [R273C] variants of p53 are present in malignant tissues [R273G] is not (27). This suggests that the different mutant-variants formed due to alterations in the same codon have different gain-of-function. In order to explore the different function of R273 variants in cancer, we have performed an analysis of The Cancer Genome Atlas. We analyzed survival probability from TCGA data (28, 29) of breast cancer patients with mutations in R273 position and WTp53 (30-32). The test estimates the Hazard Ratio (HR) using the Log-rank (Mantel-Cox) Test (33), which is the ratio between the risk of the outcome in the group possessing mutations in p53 and the group with WTp53. The HR value < 1 indicates that the WT-group has better survival than the mutant-group. As expected, we measured better survival for patients with WTp53 in breast cancer (Figure S1b, HR=0.51, 95% CI 0.28 to 0.92, p=0.026]. Similarly, HR estimations between two different types of R273 mutations indicated better survival with the [R273H]p53 mutation in Breast cancer patients (Figure S1c, HR= 0.84, 95% CI 0.3 to 2.3, p=0.74) than those with [R273C]p53. TCGA analysis confirms the variation in cancer cell malignancy among different variants of R273, where [R273C] showing minimum survival rate compared to [R273H]. These survival studies put forward a question that how R273 variants are different from each other. Till date, there is no clear understanding of the effect of R273 mutations on structure of p53. For another similar type of DNA-contact amino acid R248, reports suggest that the R248Q mutant destabilize the structure and promote the aggregation of p53 (21, 34-40). However, a meticulous correlation if these structural changes are linked to differential oncogenicity is missing in the literature. Our *in silico* studies also suggested that these variants have a different propensity for aggregation and self-assembly. We used the aggregation prediction softwares AGGRESCAN (41) and PASTA (42) to check the aggregation propensities of mutant-variants of R273 compared to the WTp53 (Figure S2). A hepta-peptide FEVXVCA has been used for the analysis, where ‘X’ is the variant amino acid. The AGGRESCAN analysis shows that the leucine variant has the maximum aggregation propensity followed by the cysteine variant. The wildtype and the histidine variant show little aggregation tendency. The PASTA analysis shows that the cysteine variant has the maximum tendency to form cross-β amyloid structures, followed by leucine, histidine and the wildtype hepta-peptides respectively. Therefore, our *in silico* prediction studies reveal that the [R273C] variant peptide is highly amyloidogenic whereas [R273L] peptide shows maximum aggregation. We next performed a spectrum of biochemical, biophysical and imaging studies to probe the difference between the self-assembly of the three p53 variants [R273L]p53, [R273H]p53 and [R273C]p53. Our studies clearly indicate stark difference between their assembly properties that might play a role in the different survival rates in different cancers. These observations reinforce the fact that personalized process specific therapeutics and medication might be a parallel pathway for cancer treatment.

**Figure1:**
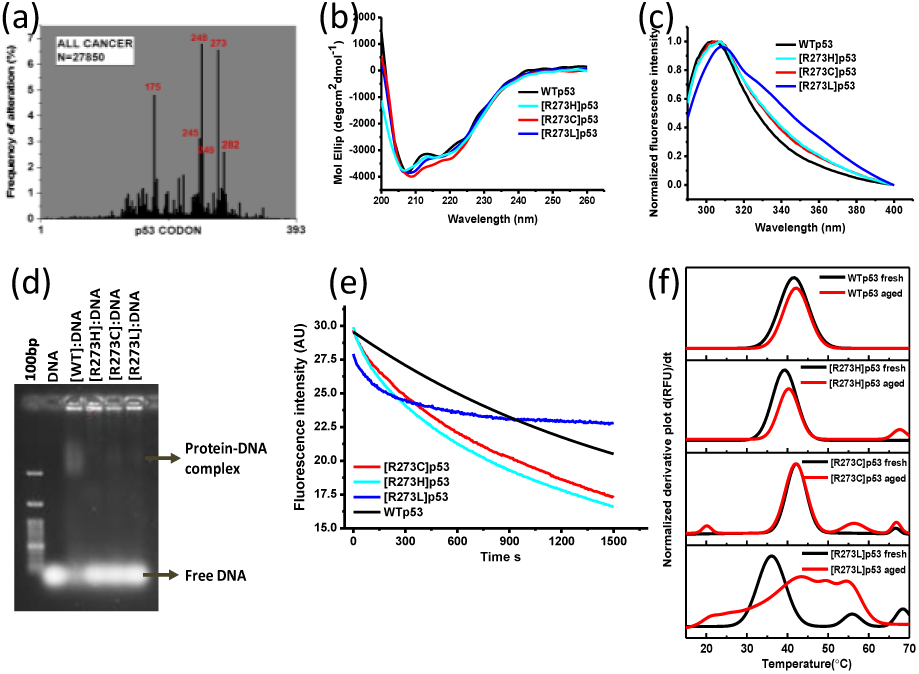
Effect of [R273]p53 variants on the stability of p53. The frequency distribution chart shows percentage of alteration of various amino acid in full length of p53 in cancer (a); The secondary structure of WTp53 (5µM) probed using circular dichroism. (inset) SDS-PAGE (12%) of purified WTp53 and its [R273X]p53protein variants (b); Intrinsic protein fluorescence of WTp53 and [R273C]p53 [R273H]p53 and [R273L]p53 mutant-variant (c); p53-DNA binding study by electrophoretic mobility shift assay shows that the DNA-binding ability is compromised in case of R273 variants compared to WT p53. (d). Change in the tyrosine fluorescence is observed at a wavelength 305 nm of WTp53 and its R273 variants (concentration of 500 nM) at 37°C as a function of time. The curves are fitted in 1storder exponential decay to calculate apparent denaturation rate(e); 1st order derivative plot obtained from thermal shift assay (TSA) of the WTp53 and its R273 variants (both fresh and aged) at a concentration of 500nM with 10X Sypro-orange. Black corresponds to fresh protein and red for aged samples of WTp53 and its variants. Thermal melt curves indicate that in case of WTp53, both aged as well as fresh protein show same transition peak whereas in case of [R273L]p53, the transition peak of unfolding vanishes and new peaks are observed in aged as compared to fresh protein. In case of [R273C]p53, the transition peak of unfolding remains same but it shows additional peaks corresponding to aggregates as well in aged sample (f).

## Materials & methods

### Chemicals

All reagents are of molecular biology grade. Ultrapure water is used for all experiments

### Propensity of aggregation of R273 hotspot variants using prediction algorithms

AGGRESCAN and PASTA2.0 algorithms are used to understand the aggregation characteristics of R273variants in hepta-peptide of p53 FEVRVCA. Aggrescan(41) is a server to predict and evaluate aggregation hotspots in polypeptide. The hotspot area of aggregation per residue is calculated. PASTA 2.0(42) (prediction of amyloid structure aggregation) predicts aggregation prone region by evaluating the stability of putative cross-beta pairings within different stretch of sequences.

### Molecular cloning and mutagenesis

The N-terminal truncated p53 (residues 82-393) construct was generously shared by C. Arrowsmith (43) Addgene (plasmid 24867). All the p53[R273] point mutations are done using overlap extension polymerase chain reaction by primers which are shown in **Table S2** and then digested with Nde1(NEB) & BamH1(NEB) and ligated in the pET15b vector. The constructs are confirmed by sequencing.

### Expression and purification of p53(82-393)

6XHis tag-p53(82-393) is expressed in *E. Coli* host strain BL21(DE3) (pLysS). Briefly, expression of p53(82-393) is induced using 0.4mM isopropyl β-D-1-thiogalactopyranoside in Luria Bertani broth (LB) containing 100 μg/mL ampicillin and 34 μg/mL chloramphenicol along with 90uM ZnSO4. After being induced at 18°C for 14 h, cells are pellet down and lysed in buffer containing 50 mM Tris-Cl(sigma-Aldrich) (pH 8.5), 200mM NaCl, 1 mg/mL lysozyme, 5% glycerol (Merck), 5 mM DTT, and 1 mM PMSF. The His6 tag p53 protein is purified from the cell lysate using Ni-NTA affinity chromatography. All the steps of purification are done at 4°C strictly.

### Intrinsic fluorescence

Intrinsic fluorescence experiments are conducted using an Edinburg Spectro-fluorometer, with excitation at 280nm and record the emission from 290nm to 400nm. A 10 mm rectangular quartz microcuvette is used for the fluorescence measurements. The change in the fluorescence intensity at 305nm is recorded with time at 37°C. Experiment is performed at a concentration of 500nM of p53 and its R273variants for 25 minutes.

### Electrophoretic mobility shift assay (EMSA)

EMSA is performed to test the DNA binding of WTp53 and its R273variants ([R273H], [R273C], [R273L])p53. The lyophilized forward primer and reverse primer of gadd45-30mer are purchased from IDT. The primers are dissolved in 10mM Tris buffer pH 8.0 to make a final concentration of 100µM. The forward and reverse primers are mixed in equal ratios to anneal them. The oligos are annealed by keeping the equimolar ratio mixture in water bath at 95°C for 10min and then slowly cool to room temperature (25°C). The protein and DNA are mixed in the ratio of 8:1(amount ratio) and an additional MgCl_2_ of 5mM. The mixture is kept at 4°C for 30min and then shifted to 22°C for 40min before loading into the 0.8% agarose gel in Tris-Borate buffer for 2hours.

### Thermal Shift Assay

TSA is a versatile technique in determining the stability of the protein. In this experiment, sypro-orange is used as a dye and the change in the sypro-orange fluorescence gives information about the unfolding of the protein. We have performed the experiment in BIORAD q-PCR machine with a protein concentration of 500nM and sypro-orange concentration of 10X at a rate of 1°C/min from 16° C till 70° C temperature.

### Circular dichroism

For measurement of CD spectra, 5µM of concentration of WT and its variants is taken. The proteins are placed into a 0.1 cm path-length quartz cell (Hellma, Forest Hills, NY). Spectra is acquired using a JASCO 810 instrument, and measurements are taken at 10°C. Spectra are recorded over the wavelength range of 200-260 nm. To study the melting temperature of a protein, the samples(5µM) are ramped from 10°C to 70°C at an interval of 5°C at a rate of 1°C/min. The molar ellipticity at 222nm as a function of temperature is observed to determine the change in helical structure of WTp53 and its R273 variants.

### Thioflavin T (ThT) Fluorescence

ThT(sigma-Aldrich) is a amyloid marker used to stain amyloids as it binds to amyloid without showing significant binding to the corresponding monomeric protein or peptide. To determine the aggregation of protein/peptide samples, a ThT binding experiment is performed. A 10mM stock solution of ThT in 100% ethanol is prepared. A working solution of 1mM of ThT is prepared in dialysis buffer. 20ul of this working solution is mixed with the protein solution(without reducing agent) such that final ThT concentration reached 100µM and the final protein concentration reached 500nM. Then the fluorescence is immediately measured. The fluorescence experiments are conducted using Edinburg fluorometer, with excitation at 440 nm and an emission range of 450-550nm. The intensity values at 490 nm are plotted.

### AFM Imaging

WTp53 and its R273 mutant-variant proteins at a concentration of 5µM are heat induced at 37°C for 1 hour. After incubation, samples are diluted at a concentration of 500nM in MilliQ water and drop-casted on silicon wafer and then washed with milliq water for 5times and dried using nitrogen gun. Samples are imaged in a soft-tapping mode in a nanoscope AFM. Sample is scanned at a scan rate of 0.5Hz. The height and width of the fibrillar aggregates are analyzed using inbuilt nanoscope analysis software.

### ANS fluorescence study

ANS binding studies are performed in Edinburg spectro-fluorometer with temperature maintained at 37°C using peltier. ANS fluorescence intensity is observed at 492nm with excitation at 375 nm with excitation and emission bandwidths set at 2 nm and 3 nm respectively. ANS emission is recorded after every 5second for 1hour at 37°C. The concentration of ANS dye is maintained at 100µM with WTp53 and its variants at concentration of 5µM.

### Ellman’s assay

WTp53 and its R273 variants fresh and aged (incubated at 37°C for 30min) are incubated with Ellman’s reagent at a molar ratio of 1:5 (1µM of protein with 5µM of Ellman’s reagent) for 5min and then take absorbance in UV-visible spectrometer from 200nm to 600nm.

### Cytotoxicity using MTT assay

Relative cell viability is determined using MTT (sigma-Aldrich) assay. HEK293T cells (10^4^ cells/well) are seeded in a 96-well plate. After 12 hours incubation, the cells are replenished with media containing different conc. of proteins (300nM to 1µM) for 24 hrs. Cells treated with medium only served as a negative control group. Similarly, MCF-7 cells are seeded in a 96-well plate and after 12 hours of incubation, WTp53 and its R273 mutant plasmids (mammalian) are transfected in the concentration range of 0.5-4µg/µl using lipofectamine (Thermo-fisher). After 24 h, 10 μl of MTT solution (5 mg ml in PBS) is added to each well. After incubation for another 4 hours, the resultant formazan crystals are dissolved in DMSO (100μl) and the absorbance intensity measured by a microplate reader (Infinite M plex 200, Tecan) at 580 nm with a reference wavelength of 630 nm. All experiments are performed in triplicates, and the relative cell viability (%) was expressed as a percentage relative to the untreated control cells.

### Molecular Dynamics Simulations

All the molecular dynamics (MD) simulations were setup using the QwikMD(44) plugin of VMD(45) and run using NAMD(46) version 2.12. Short initial peptide structure are obtained from WTp53 (PDB ID: 1TUP). Then the peptide structure is energy minimized first in vacuum and then in solvated state. All the mutation has been performed in-silico using QwikMD only. The mutated structure is also energy minimized using the same protocol. The energy minimized structure is then subjected for 20 ns long MD simulation. Then MD trajectory is analyzed using grcarma. The most populated conformation obtained from MD simulation is used as the initial structure for the aggregation simulation. Then 25 peptides are arranged in random orientation but equidistant form each other having distance of 40Å from its neighboring peptide in a square shaped fashion. After the addition of hydrogen atoms, the structures are aligned with the longest axis along the Z-axis and solvated with TIP3P(47) water to form a box with a distance of 15Å between the edge of the box and the protein in all the axis. The Na^+^ and Cl^-^ ions corresponding to a concentration of 150 mM are then placed randomly in the water box by replacing the water molecules. The system is then minimized for 5000 steps with the position of the protein atoms fixed followed by 5000 steps without any restraints. The temperature of the system is then gradually raised to 300K at a rate of 1 K every 600 steps with the backbone atoms restrained. Thisis followed by equilibration for 5 ns with the backbone atoms restrained followed by 10 ns of the production run with no restraints. For all the steps, the pressure is maintained at 1 atm using Nose-Hoover method(48), thelong-range interactions were treated using the particle-mesh Ewald (PME)(49) method and the equations of motion are integrated using the r-RESPA scheme to update short-range van der Waals interactions every step and long-range electrostatic interactions every 2 steps.

## Results

### Expression and Purification of WT and R273 variants of p53

Four R273 variants of p53 viz. WTp53, [R273L]p53, [R273H]p53 and [R273C]p53 are expressed in bacterial *E.coli* with the N-terminal truncated sequence as already reported and purified using Ni-NTA affinity chromatography (43). The purification of full length p53 lead to challenges including protein aggregation and misfolding during purification, hence the N-terminal truncated form is used here. We monitored the structure and the stability of all the variants using circular dichroism and intrinsic protein fluorescence (Figure 1b & 1c). All the mutant-variants except the [R273L]p53 showed fluorescence spectrum similar to the WTp53. The mutant [R273L]p53 shows some initial destabilization as observed by the broadening of the fluorescence spectrum compared to the WTp53 at 4°C. However, the circular dichroism spectra are similar in all variants suggesting no significant alteration in the secondary structure (Figure 1b).

### Investigating the DNA-binding of R273 variants of p53

The R273 is one of the DNA-contact positions in p53 that allows specific binding between p53 and DNA. The mutation of this amino acid to any other amino acid leads to altered DNA binding. To test this hypothesis, we have employed an electrophoretic mobility shift assay (EMSA). EMSA is an effective way to screen for the DNA-binding characteristic of proteins (50-52). We have selected the Gadd 45-30mer dsDNA (Table S2) due to its specific binding capability with WTp53(53). As the c-terminal of p53 is reported to bind non-specifically with DNA(54,55), in all our experiments we have used a salt concentration of 200mM in Tris-Cl buffer of pH 8.5 and 7.5 respectively is used for the assay to avoid such non-specific binding. At a pH of 7.5, non-specific binding is there due to electrostatic interactions between negatively charged DNA and positively charged c-terminal as shown in Figure S3. On the other hand, at a pH of 8.5, the net positive charge on protein decreases resulting in a decrease in non-specific binding. WTp53 and the [R273X]p53 variants are incubated at a ratio of 8:1 with dsDNA (8µg of protein with 1µg of dsDNA) at 4°C for 30min then shift to 22°C for 40 min followed by agarose gel electrophoresis in 1X TB buffer. At pH 8.5, we observe retardation in the mobility of DNA incubated with WTp53 in contrast to when incubated with the R273 mutant-variants. This indicates that only WTp53 binds to DNA and the mutants cannot (Figure 1d). Also we observe, free DNA in the lanes with of R273 mutant-variants whereas p53-DNA complex in the lane withWTp53. Following the establishment of the loss of function due to mutation, we next aimed to study if there are structural implications as well due to these mutations in addition to the functional.

### Stability studies of p53 R273 mutant-variants in vitro

It has been reported earlier that the native folded structure of p53 shows predominantly tyrosine fluorescence whereas tryptophan fluorescence prevails in case of denatured p53(56). Post purification (4°C) except for the [R273L]p53 variant all other variants show tyrosine fluorescence similar to the WTp53. The [R273L]p53 show slightly broadened spectra indicating alterations in the positions of the tyrosine/tryptophan residues. Next, we shifted all the variants to the physiological temperature of 37°C. Under this condition, we observe a change in the tyrosine fluorescence for all the variants compared to the WTp53. To compare the stability of R273 variants with that of WTp53, we have performed studies on the time dependent change in tyrosine fluorescence. While in the case of the WTp53, a gradual decrease in the fluorescence intensity at 305 nm is seen, all the other variants show a faster decrease (Figure 1e). The decay plots are fitted to the following equation *y* = *Ae*^-*kx*^ + *y*0 to calculate the apparent rate (*k*) of exposure of tyrosine residues which can be related to the denaturation of the protein. The WTp53 has an apparent denaturation rate (*k*_*wt*_) of 0.6 × 10^-3^±1.1×10^-5^s^-1^while those for the [R273H]p53, [R273C]p53, [R273L]p53 variants are 1.3×10^-3^±1.5×10^-5^s^-1^,1.1×10^-3^±1.4×10^-5^s^-1^ and 3.2×10^-3^±4.5×10^-5^s^-1^ respectively (Figure 1e) indicating that [R273L]p53 denatures 5 times faster than WTp53 whereas [R273C]p53 and [R273H]p53 variants unfold 2 times faster than WTp53.

We next assessed the thermal stability of the variants using thermal shift assay (TSA) probed by sypro-orange fluorescence (Figure 1f, Figure S4). TSA of the fresh WTp53 shows only one transition around 40°C. [R273H]p53 also shows a single transition in the same region. However, the variants [R273C]p53 and [R273L]p53 in addition to the transitions around 40°C show additional transition centered around 60-70°C. Next, we allowed the variants to age for 1 hour and performed TSA. The aged WTp53 and [R273H]p53 behave similar to that of the fresh samples with one transition at 40°C. In the case of [R273C]p53 aged samples we observe additional higher temperature transitions in addition to the one observed in the fresh samples and still show a transition similar to the fresh samples. For the [R273L]p53a completely different transition curve is observed. Unlike the other variants, that start to denature ∼40°C, however, the aged [R273L]p53 starts the transition at a much lower temperature of 20°C and several transitions including some very high temperature transitions are observed. The *T*_*m*_ of the fresh samples obtained from the TSA and CD melts are shown in (Table S3, Figure S4, and S5). We see that the *T*_*m*_ of the variants is lower than that of the wild type form. These studies indicate that the mutation at the R273 position destabilizes the structure of the protein leading to altered structure and folding. In the next set of studies, we focused on understanding if the loss in the stability of the variants lead to misfolding and aggregation as observed for some other p53 variants in the literature.

### Amyloidogenic behavior of R273 variants of p53

We studied the aggregation characteristics of the R273 variants using Thioflavin T (ThT) as reported in the literature (7, 20, 38, 40, 57, 58). All the variants are incubated in Tris buffer at pH of 8.5 containing 200mM NaCl and 100 µM ThT and the interaction with ThT is studied by exciting the mixture at 440 nm and recording the fluorescence intensity at 490nm.We observed that the initial binding with ThT is maximum for the [R273C]p53 while negligible interaction is observed with all other variants (Figure 2a). Incubation of the variants at 37°C leads to enhanced ThT fluorescence with time indicating formation of amyloid like aggregates. The enhancement is maximum, in the case of [R273C]p53(although in the presence of DTT, there is no significant change in the ThT binding of WTp53 and [R273C]p53) followed by [R273L]p53, while the [R273H]p53 and WTp53 show minimal change (Figure 2b). The [R273C] variant also shows the maximum Congo red dye absorbance followed by [R273L]p53. The [R273H] and WTp53 show the least interaction with the dye (Figure 2c).

**Figure 2:**
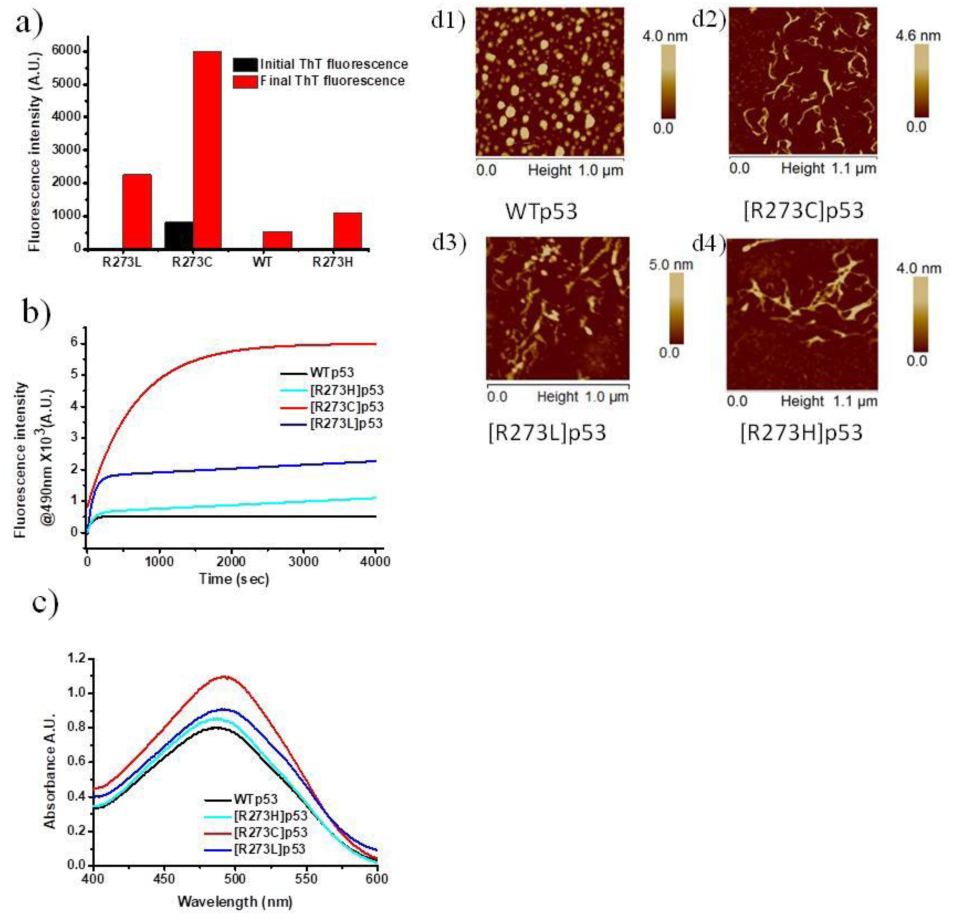
Amyloidogenicity of [R273X]p53 protein variants. Bar Graph of initial and final values of fluorescence intensity of ThT incubated with WTp53 and its variants (a); Fluorescence intensity of thioflavin T incubated with WTp53 and its R273 variants. The thioflavin T fluorescence of [R273C]p53 increases with time at 37°C but it get saturated for other variants and WTp53(b); Congo red absorbance for the aged WTp53 and variants. It is observed that [R273C]p53 shows maximum absorbance than another any other variant (c); AFM images of WTp53 and its R273variants show the globular and fibrillar assemblies respectively (d (1-4));

These studies indicate that the aggregates of the R273 variants are amyloidogenic in nature and the amyloidogenicity follows the order [R273C]p53>[R273L]p53>[R273H]p53≈WTp53. These results are corroborated with *in silico* aggregation prediction studies. We next checked the morphology of the aggregates post ThT studies. For this, an experiment is designed where the aggregation was initiated in Tris buffer at pH of 8.5 containing 200mM NaCl without ThT. After 1hr of incubation, the samples are drop-casted on a silicon wafer and imaged using AFM (Figure 2d1-d4). The wild type protein shows globular assembly, while all the oncogenic variants show fibrillar assemblies typical to those of amyloid fibres. Although [R273H]p53 shows sparsely populated fibrillar aggregates. This is exactly in accordance to what we have observed in our experiments involving ThT and Congo red binding.

### Understanding the molecular mechanism for the unique stability and self-assembly of different R273 variants of p53

Next, we attempted to understand the mechanism of aggregation in the different variants that leads to the altered self-assembly. We used the ANS fluorescence studies to check how the different variants of the protein unfold during the aggregation process. ANS is a dye that binds to the solvent exposed hydrophobic patches of protein. In our experiments, we see that except for the wild type in the case of all the three variants, there is a monotonous increase in the ANS fluorescence with time immediately after the incubation at 37°C. The WTp53 shows a lag phase of close to 10 minutes before the exposure of the hydrophobic patches. For the [R273C]p53 variant, after an initial increase, there is a sharp drop in the ANS fluorescence indicating initial destabilization of the protein by exposure of hydrophobic surface followed by formation of intermediate structures with buried hydrophobic surfaces (Figure S6). We next performed the estimation of free thiol groups using the Ellman’s reagent. For the WT, [R273L]p53 and [R273H]p53 variant, we observe that there is no spectral shift when the unaggregated and the aggregated samples are reacted with the reagent (Figure S7). On the other hand, for the [R273C]p53 protein, there is a marked difference in the interaction of the unaggregated sample with that of the aggregated one. Qualitatively, the number of free thiol groups in the case of the aggregated samples is less than that of the unaggregated ones. This suggests that in the case of [R273C]p53, the aggregation is disulphide mediated unlike the other variants used in this study. Further, we have done maleimide linked dye labelling experiment to understand the status of disulphide bond in WTp53 and [R273C]p53. For this, we have overnight incubated WTp53 and [R273C]p53 with cy5 maleimide in the ratio of 1:5 at both 4°C and room temperature (Figure S8). The free dye is removed and then check the concentration of the dye bound protein and perform cy5 fluorescence studies. We excited the dye bound protein at 640 nm and take the emission from 660 nm till 750 nm. At a same concentration of WTp53 and its [R273C]p53 variant, the cy5 fluorescence intensity of WTp53 is almost same for 4°C and room temperature condition whereas it gets decrease by around 30% in case of [R273C]p53 variant. These experiments support the hypothesis that [R273C]p53 self-assembly is mediated by disulphide bond as shown in figure S9. We have also verified the disulphide bond partners in WTp53 and [R273C]p53 using DIANNA server (Figure S9) (59,60).

### Molecular dynamics simulations

We next probed the aggregation of p53 variant proteins in-silico using all-atom MD simulations. The 15-residuepeptide stretch of ^262^GNLLGRNSFEV<u>X</u>VCA^276^in explicit water (TIP3P) in 150 mM KCl is monitored for 60 ns. The inter-peptide separation at the beginning of the simulation is set to 40 Å to avoid non-specific contacts during free rotations of the peptides. The peptide stretch is one complete β–strand that includes the 273 position (here X). Figure 3 shows the snapshots of the peptides at various time points of the simulation for WTp53^262-76^ (a), [R273H]p53^262-76^ (c), [R273C]p53^262-76^ (e), [R273L]p53^262-76^ (g). When monitored the inter-peptide H-bonds with time (Figure S10), we notice the early on-set of self-assembly for the oncogenic variant peptides where the cluster size increases aggressively with time (Figure 3). [R273L]p53 formed the largest cluster at the end of the simulation. In contrast, WTp53^262-76^ peptides form small clusters which dynamically dissociate and associate throughout the simulation. We next plotted the minimum distance between all pairs of the peptides in the last frame of the simulation as a contour to get a quantitative idea about the contacts and inter-chain separation of the peptides. The inter-peptide distances for the WTp53 are substantially higher as expected compared to all the oncogenic variant peptides. For the WT, a greater number of peptide pairs that reside in the range of 20-40 nm while for all the other peptides, the peptide pairs are 0-20 nm apart.

**Figure 3:**
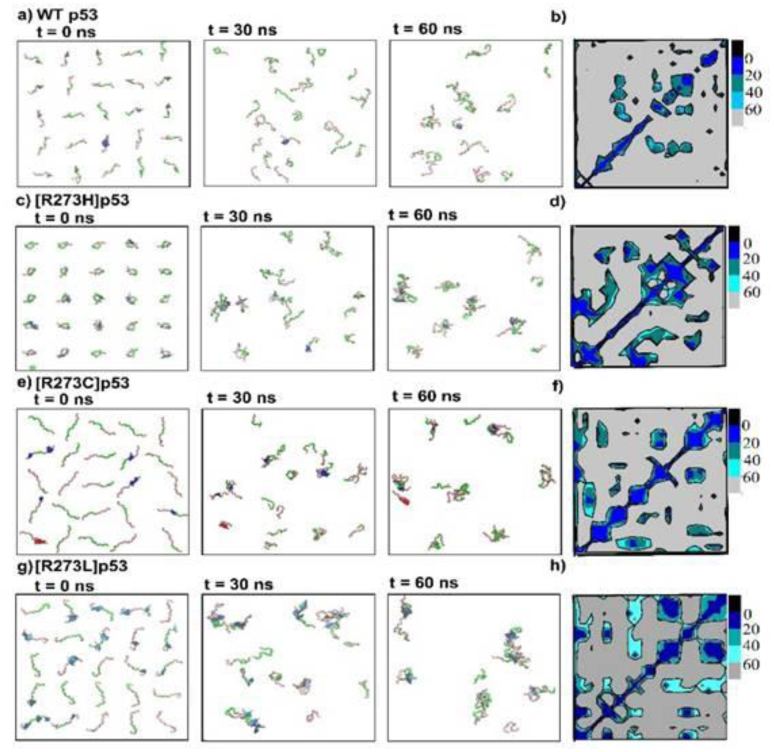
Molecular dynamics simulation of ^262^GNLLGRNSFEVHVCA^276^peptide and its different variants at R273 position. Snapshots at various time points during the simulation (0 ns, 30 ns and 60 ns) for WTp53^262-76^ peptide (a); [R273H]p53^262-76^ (c); [R273C]p53^262-76^ (e);[R273L]p53^262-76^ (g). The right panel shows the corresponding distance map of the peptides at the last frame of the simulation (b, d, f, h). Distance has been calculated between the geometric center of corresponding two peptides. Black represents 0 nm distance, blue represents 0-20 nm distance, deep cyan represents 20-40 nm, light cyan represents 40-60 nm and gray represents all the inter-peptide distances above 60 nm.

Interestingly, the inter-residue contact-maps from the peptides feature different patterns of interaction for different variants of the peptide-stretch indicating that the initial nuclei formation is not same. Similar inferences are also made from the ThT kinetics, ANS binding and the studies using the Elman’s reagent. Further, we observed that the development of the secondary structures during the simulation is different for the different variants (Figure 4). The peptide-stretch resembling WTp53 mostly remain as random-coil. In fact, only two out of twenty-five peptides showed random coil to helix, and five peptides showed random coil to transient β-strand transformation (Figure 4a). In contrast, appearance of the secondary structures is frequently observed in all the variant peptides (Figure 4b,c,d). All the variants undergo secondary structural transition to β-strand very frequently. [R273L]p53 shows the greatest extent of β-strand transformation (Figure 4d). Notably, [R273C] also undergo transition to α-helical structure and to pi-helix in a greater extent which may be one of the key to its different aggregation propensity. As for tau protein involved in Alzheimer’s disease, α-helix is considered to initiate the aggregation process(61).These structural transformations could be the key to stabilize as well as increase the size of the aggregates for mutant-variants, as clusters with parallel and anti-parallel β-strand organization are also noticed in the simulations.The difference in the contact map, H-bonds and dissimilar secondary structure formation in the course of the simulation suggest that the proteins misfold via different aggregation pathways, which in turn can result in different levels of pathogenic aggression as observed in the TCGA analyses.

**Figure 4:**
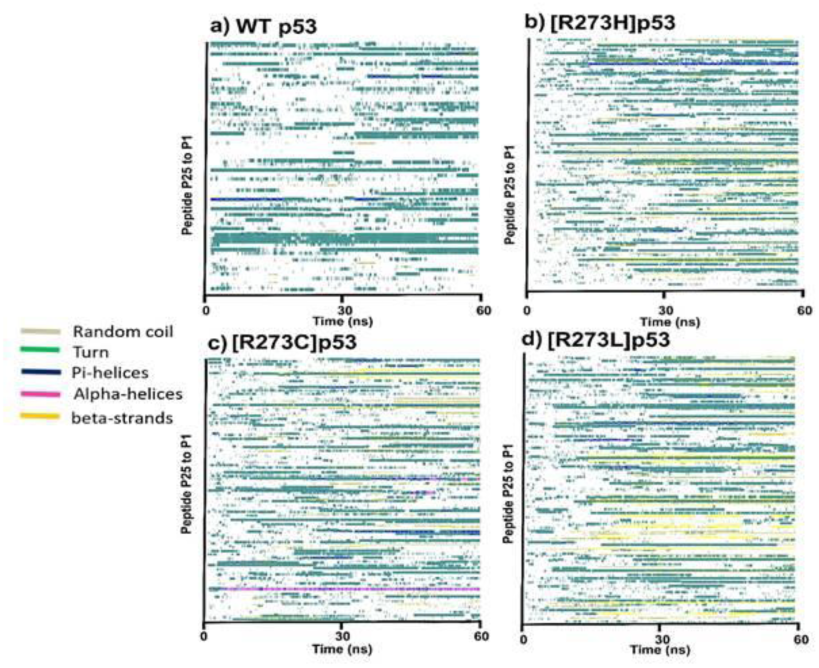
Change in the secondary structure of various peptides from P1 to P25 with the progress of the simulation from 0 ns to 60 ns. WTp53 (a); [R273H]p53(b); [R273C]p53(c); [R273L]p53(d). Various color like grey, green, blue, pink, and yellow represent random coil, turn, π-helices, α-helices, and β-strand, respectively.

### Cytotoxicity mediated by the [R273X]p53 oligomers

It is a well-documented fact that amyloidogenic oligomers of several proteins including p53 are toxic and lead to reduced cell viability (62,63). We hence assessed the effect of the oligomers of R273 variants on the viability of HEK293 cells using MTT assay. The variant concentration is used in the range of 30nM−1µM. Our experiments show that the WTp53 is not toxic in the said concentration range. The [R273L]p53 and [R273C]p53 variant start showing toxic effects at a concentration as low as 30 nM and show close to 50% cell viability at 300 nM concentration. Increasing the concentration to 1µM did not show any further decrease in the viability. [R273H]p53 shows similar cell viability as the WT till 300 nM concentration and ∼80% cell viability at 1 µM concentration as shown in Figure 5a. Our toxicity data is in support of our biophysical studies, where we show that the different variants have different stability, modes of denaturation and aggregation propensities. We next checked the cell viability in breast cancer cell line MCF-7 cells by the*in situ* over-expression of WTp53 and the various [R273X]p53 variants. We see that the cell viability decreases in the presence of WTp53 while all the [R273X]p53mutant-variants enhance the cell survival of MCF-7 (Figure 5b). We have already observed in the TCGA analysis for the cancer patients that the R273 variants decrease the survival of the patients (Figure 5b). This decrease in the survival may be because of the enhanced viability conferred to the cancer cells by the mutant-variants as seen in these experiments.

**Figure 5.**
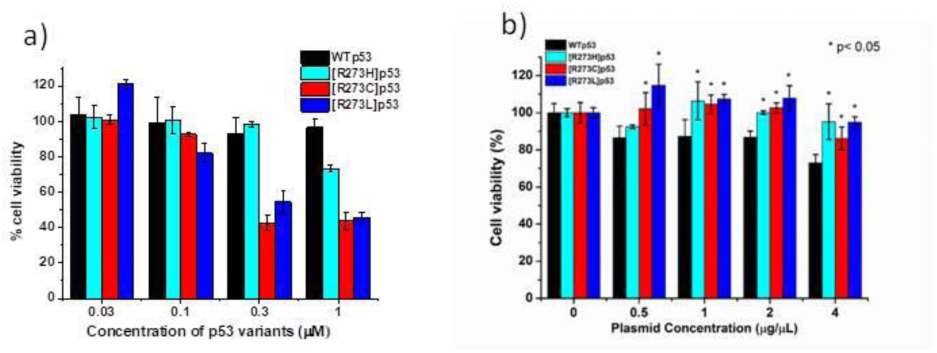
Toxicity of p53 variants. The cellular toxicity induced by the p53 (82-393) WTp53 protein and its variants is determined by using MTT assay. The HEK293T cells are treated with WTp53 and its R273 variant proteins in the concentration range from 30 nM-1 µM for 12 hrs. The percentage cell viability of [R273C]p53 is minimum followed by [R273L]p53 and [R273H]p53. WT p53 shows almost 100% viability in this concentration range (a). Cell viability assessment of over-expressed full length WTp53 and its R273 mutant-variants in MCF-7 cell line. Different concentrations (0.5µg/ml to 4µg/ml) of plasmids are transfected in MCF-7 cell line and incubated for 24h before treating them with MTT. Cell viability decreases in the presence of WTp53 whereas more cells are viable in case of R273 variants. All the variants show significant viability compared to the WT (b).

## Discussion

Mutations in physiological proteins lead to several pathogenic conditions and p53 is one such classic example. Being central to several physiological processes, a slight anomaly in its genetic makeup leads to clinical conditions. In this study, we selected the R273 position in p53 to study the effect of the pathological R273 mutant-variants on the structure and function of the protein. The R273 position is a DNA contact position and interestingly mutated to different amino acids leading to different forms of cancer. The range of mutations causes different extent of pathogenic aggression. Our studies show that each of the variant proteins are unique in their ways when it comes to the structure-function relationship. All the variants, unlike the WT, show compensated DNA binding. This result is apparent as the position R273 is a DNA contact amino acid and hence, changing it to any other residue will lead to loss of the function. However, as we argued earlier, if lack of DNA binding is the only reason for cancer pathogenesis, then why the different variants will lead to different extent of clinical symptoms and side effects. This can happen only if there are associated structural and functional roles of these oncogenic variants in physiology.

All the three mutant-variants studied show altered stability compared to the WTp53. The [R273L]p53 variant show maximum instability. This instability can be because[R273L]p53mutation is from a charged residue to a non-conservative hydrophobic residue. This can cause to local changes in the structure leading to instability as seen in our experiments. Our experiments indicate broadening of the protein fluorescence spectra indicating some modification in the structure compared to the wild type protein. However, in the absence of the crystal structure we cannot comment exactly on the extent of the local structural alterations. Corroborating with our experiments, the simulation studies also show maximum propensity of this variant towards formation of aggregation clusters early on, in contrast to the WTp53 peptide. The crystal structures of the [R273C]p53 and [R273H]p53 are similar to those of the wild type protein but it was solved in the presence of four stabilizing mutations (64,65). Our experimental and aggregation prediction server studies reveal that [R273C]p53 is highly amyloidogenic among these variants. Our biochemical studies and disulphide prediction(59) shows that [R723C]p53 forms shuffled disulphide bonds indicating that [R273C]p53 aggregation may be mediated by disulphide bonds. The [R273C]p53 most prevalent in the cancer of the brain where the condition is oxidizing (66,67). Also, the role of disulphide bonds in the self-assembly and amyloidogenicity of neurodegenerative diseases is well documented in the literature (68). Hence, our studies indicate that understanding the link between disulphide mediated amyloid aggregate formations and brain cancer will be an interesting area to explore in future.

## Conclusion

Our studies on the different R273 variants of p53 clearly indicate the role of individual amino acids in the stability and self-assembly of the protein. It is interesting to note that changing a single amino acid leads to different structure-function relationship in the protein. These studies also emphasize that it is not only the lack in DNA contact that leads to p53 inactivation in certain cancers, but also the structural changes that are associated with the mutations. These observations, thus, call for the need of developing therapeutic strategies that will consider the hallmark of each of these mutations as a therapeutic target rather than considering the mutation as the target. We do understand that there may be other simultaneous mutations in the *p53* gene that also affect the pathogenesis of the disease, however, our studies indicate that specific therapeutic strategies that precisely target the additional effect of the mutation apart from compromised DNA binding is also required for cancer management and cure.

## Abbreviations

AFM: Atomic Force Microscopy;
Ds: Double stranded;
EMSA: Electrophoretic Mobility Shift Assay;
WT: Wild Type;
TSA: Thermal Shift Assay;
ThT: Thioflavin T;
TCGA: The Cancer Genome Atlas.

## Author contributions

SS and SR designed the research. AG, MKS, JPH conducted all the experiments. AG, JPH, SS and SR analyzed and interpreted the data. AG, SS and SR wrote the manuscript.

## Acknowledgements

The authors acknowledge the members of SS and SR lab for critical insights in the manuscript. The authors acknowledge Ms. Simerpreet Kaur for help in designing certain experiments. The study supported from the institutional seed fund to SS and SR. The authors declare no conflict of interest.

## TOC Graphics

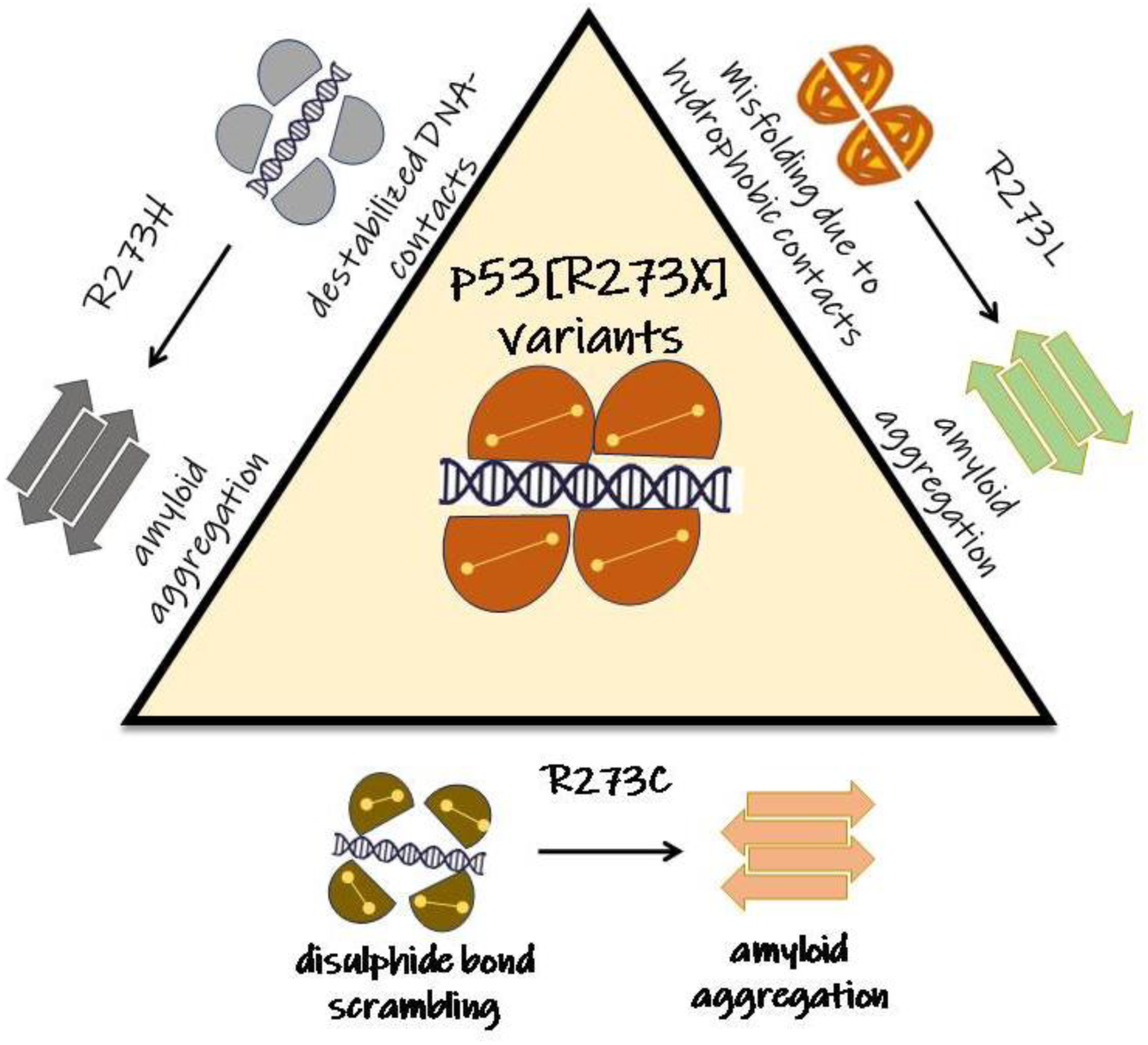

